# An updated model of shoot apical meristem regulation by ERECTA family and CLAVATA3 signaling pathways

**DOI:** 10.1101/2023.09.29.560237

**Authors:** Muhammad Uzair, Ricardo Andres Urquidi Camacho, Ziyi Liu, Alex M. Overholt, Daniel DeGennaro, Liang Zhang, Brittani S. Herron, Tian Hong, Elena D. Shpak

**Affiliations:** Department of Biochemistry, Cellular and Molecular Biology, University of Tennessee, Knoxville, TN, 37996, USA; UT-ORNL Graduate School of Genome Science and Technology, The University of Tennessee, Knoxville, Tennessee, USA; Complex Carbohydrate Research Center, University of Georgia, Athens, GA 30602, USA

**Keywords:** Arabidopsis, stem cells, ERECTA, WUSCHEL, CLAVATA3, shoot apical meristem

## Abstract

The shoot apical meristem (SAM) gives rise to above-ground organs. The size of the SAM is relatively constant due to the balance of stem cell replenishment versus cell recruitment into developing organs. In angiosperms, the transcription factor WUSCHEL (WUS) promotes stem cell identity in the central zone of the SAM. WUS forms a negative feedback loop with a signaling pathway activated by CLAVATA3 (CLV3). In the periphery of the SAM, the ERECTA family (ERf) receptors promote cell differentiation and constrain the expression of *WUS* and *CLV3*. Here, we show that four ligands of ERfs redundantly inhibit *CLV3* and *WUS* expression. Transcriptome analysis confirmed that *WUS* and *CLV3* are the main targets of ERf signaling and uncovered several new ones. Analysis of promoter reporters indicated that in the vegetative meristem, the *WUS* expression domain mostly overlapped with the *CLV3* domain and did not shift along the apical-basal axis in *clv3*. A 3D mathematical model reproduced the experimentally observed *CLV3* and *WUS* expression patterns with fewer assumptions than earlier models. Based on these findings, we propose that CLV3 regulates cellular levels of *WUS* expression through autocrine signaling, while ERfs regulate *WUS* spatial expression, preventing its encroachment into the peripheral zone.

**One Sentence Summary:** Through autocrine signaling, CLV3 regulates the level of *WUS* expression in the vegetative SAM but not its location, while ERfs regulate the *WUS* spatial pattern, preventing its expansion into the peripheral zone.

## INTRODUCTION

Plant meristems contain a pool of undifferentiated cells that are used for organogenesis throughout the life of an organism. The shoot apical meristem (SAM) forms between cotyledons during embryogenesis. After germination, it generates the main stem’s internodes, leaves, and flowers. Later in development, axillary meristems develop in the leaf axils and form branches. In all meristems, there is a small cluster of pluripotent stem cells in the center. Once cells are displaced from the center into the periphery, they grow and divide at a faster rate, differentiate, and ultimately are incorporated into organs. The molecular mechanisms controlling the transition of stem cells into differentiating cells of the peripheral zone are of fundamental interest to plant developmental biology. In the SAM, this transition strongly relies on the ability of cells to communicate using extracellular signals.

The homeobox transcription factor WUSCHEL (WUS, AT2G17950) is essential for maintaining the SAM’s central zone in Arabidopsis. In the *wus* mutant, cells in the center differentiate prematurely, and the SAM disappears (Laux et al., 1996; Mayer et al., 1998). Ectopic and inducible expression of *WUS* promotes stem cell identity and increases the size of the central zone (Schoof et al., 2000; Yadav et al., 2010). Multiple signaling pathways regulate the expression of *WUS*. Cytokinins promote and position *WUS* expression along the apical-basal axis (Lindsay et al., 2006; Gordon et al., 2009; Chickarmane et al., 2012). A signaling pathway activated by the extracellular glycopeptide CLAVATA3 (CLV3, AT2G27250) inhibits *WUS* expression. When CLV3 or its putative receptor CLAVATA1 (CLV1, AT1G75820) are mutated, expression of *WUS* is increased (Clark et al., 1993, 1995; Brand et al., 2000; Schoof et al., 2000). In turn, WUS promotes the expression of *CLV3,* which leads to the formation of a negative feedback loop responsible for the stability of the SAM size (Schoof et al., 2000).

Recently, we demonstrated that in addition to CLV3, another signaling pathway controls *WUS* expression (Zhang et al., 2021). In Arabidopsis, three plasma-membrane receptors, ERECTA (ER; AT2G26330), ERECTA-LIKE 1 (ERL1; AT5G62230), and ERL2 (AT5G07180), redundantly regulate the width of the vegetative SAM and promote leaf initiation in its periphery (Chen et al., 2013; Uchida et al., 2013). Collectively, these receptors are called ERfs (for **ER**ECTA **f**amily receptor**s)**. In the SAM, ERf activity is controlled by four extracellular proteins EPFL1 (AT5G10310), EPFL2 (AT4G37810), EPFL4 (AT4G14723), and EPFL6 (AT2G30370) that are expressed in the SAM’s periphery (Kosentka et al., 2019). Genetic analysis demonstrated that ERfs and CLV3 function synergistically in controlling the SAM size and organogenesis in the peripheral zone (Kimura et al., 2018; Zhang et al., 2021). The *clv3 er erl1 erl2* mutant forms a gigantic meristem that cannot form leaves or internodes (Zhang et al., 2021). Our data showed that *wus* is epistatic to *ERf* genes (Zhang et al., 2021). Stimulation of ERf signaling with exogenous EPFL4 or EPFL6 rapidly decreased both *CLV3* and *WUS* expression. Based on these data, we proposed that ERfs restrict the width of the central zone in the SAM by inhibiting the expression of *CLV3* and *WUS* in the peripheral zone (Zhang et al., 2021).

Here, we investigated the role of EPFLs in the regulation of *CLV3* and *WUS* expression and uncovered that four EPFL ligands function redundantly. Using RNAseq, we analyzed gene expression changes after a brief activation of ERf signaling with EPFL6. This experiment confirmed that *CLV3* and *WUS* are the main targets of the pathway and uncovered several new potential targets. In addition, we studied the role of CLV3 in the control of *WUS* expression.

While it is broadly accepted that CLV3 prevents *WUS* expression in the top layers of the meristem, our analysis indicated that CLV3 regulates the amount of *WUS* per cell and not its spatial expression. Finally, the role of EPFL signaling in leaf organogenesis was studied using the DRN and DRNL markers.

## RESULTS

### EPFL1, EPFL2, EPFL4, and EPFL6 redundantly control expression of WUS and CLV3

Four EPFLs (EPFL1, EPFL2, EPFL4, and EPFL6) redundantly restrict the size of the SAM (Kosentka et al., 2019). When EPFL4 and EPFL6 are supplied exogenously, they suppress *WUS* and *CLV3* (Zhang et al., 2021). To test whether EPFL1 and EPFL2 also regulate the expression of *WUS* and *CLV3*, we analyzed the spatial expression of *CLV3* and *WUS* in the vegetative SAM of *epfl* mutants using previously described H2B-GFP reporters (Zhang et al., 2021). Three days post germination (DPG) seedlings were used for all experiments. The reporter analysis showed that simultaneous knockout of either *EPFL4* and *EPFL6* or *EPFL1* and *EPFL2* had a minimal effect on the spatial expression pattern of *CLV3* and *WUS* (Fig. 1A-D). We observed only a minute increase in the height of *WUS* domain in *epfl1,2* and *epfl4,6* mutants (Fig. 1A and B) and a very slight broadening of *CLV3* in the L1 layer of *epfl1,2* (Fig. 1C and D). The small increase in *CLV3* expression in the *epfl1,2* mutant correlated with a slightly broader SAM (Fig.1E). Next, we analyzed the expression of *CLV3* and *WUS* in *epfl1/+,2,4,*6 and *epfl1,2,4,6* seedlings. In seedlings heterozygous for *epfl1* mutations, the sizes of *CLV3* and *WUS* domains were slightly increased compared to both the wild type and double mutants (Fig. 1A-D). Again, this correlated with a subtle increase in the SAM width (Fig. 1E). In the quadruple *epfl1,2,4,6* mutant, the width of *WUS* and *CLV3* expression domains and the SAM were the most significantly increased (Fig. 1A-E). These experiments indicate that all four ligands regulate the expression of *WUS* and *CLV3* in a mostly redundant manner.

**Figure 1.**
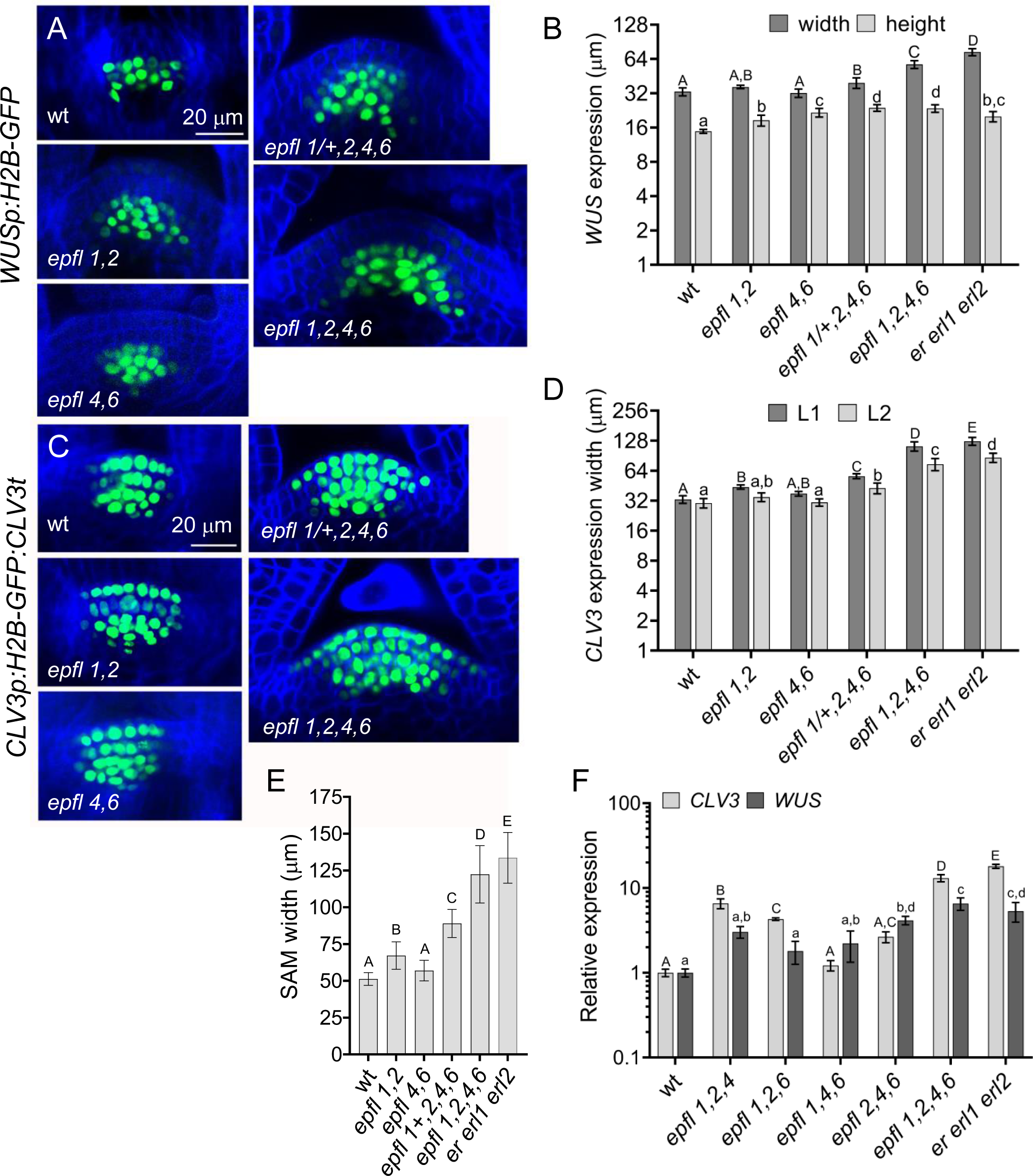
*EPFL* genes synergistically regulate the expression of *CLV3* and *WUS*. A and C. Representative confocal images of the SAM region of 3 DPG wild-type (wt) and mutant seedlings transformed with *WUSp: H2B-GFP* or *CLV3p:H2B-GFP: CLV3t* (green). In a panel, all images are under the same magnification. The cell walls were stained with SR2200 (blue). B. The width and height of *WUS* expression. n=7-20. D. The width of *CLV3* expression in the L1 and L2 layers. n=8-15. E. Comparison of the SAM width at 3 DPG in the wt, *er erl1 erl2*, and *epfl* family mutants. n=13-54. B,D, and E. Measurements were done using confocal images. F. RT-qPCR analysis of *WUS* and *CLV3* in 3 DPG seedlings. ACTIN2 was used as an internal control. B, D-F. Data is average ± S.D. Statistical differences were detected using one-way ANOVA followed by Tukey post-hoc test with a minimum *P-*value of <0.05.

At the same time, there are subtleties in the contribution of individual EPFLs to the regulation of these two genes. When we analyzed the expression of *WUS* and *CLV3* in *epfl* triple and quadruple mutants using RT-qPCR, we observed increased expression of these two genes, and the mutant combination dictated which one was upregulated more (Fig. 1F). The experiment suggests that EPFL1 and EPFL2 might have a more significant role in the regulation of *CLV3* while EPFL4 and EPFL6 are more important for the regulation of *WUS*. In addition, a comparison of *epfl1,2,4,6* and *er erl1 erl2* mutants identified some differences. In *epfl1,2,4,6, CLV3* and *WUS* were expressed in a narrower domain (Fig. 1B and D), *CLV3* was expressed at a lower level (Fig. 1F), and the SAM was slightly narrower (Fig. 1E). This suggests that either ERf receptors can weakly regulate the SAM in a ligand-independent manner or additional EPFL ligand(s) contribute(s) to the regulation of the SAM structure.

### Transcriptome analysis identified several meristematic genes, including *CLV3* and *WUS*, as the immediate targets of EPFL6

Transient activation of ERf receptors with EPFL4 and EPFL6 followed by RT-qPCR identified *CLV3* and *WUS* as downstream targets (Zhang et al., 2021). To discover targets of the ERf signaling using an unbiased approach, we performed transcriptome sequencing (RNA-seq). As we were specifically interested in the meristematic targets, many of which are expressed at low levels, we used *clv3 epfl1 epfl2 eplf4 epfl6* seedlings that have very large vegetative SAMs (Zhang et al., 2021) and performed a relatively deep sequencing (>50M reads per sample). We treated three DPG seedlings exogenously with 10 μM EPFL6 for 3 hours with and without 10 μM cycloheximide (CHX). Cycloheximide was used to test whether the regulation of gene expression by EPFL6 depends on translation. Based on principal component analysis (PCA), the obtained RNA-seq data clustered according to treatment and showed a high degree of intra-treatment reproducibility (Supplemental Fig. 1A). Pearson correlation heatmap of replicates also indicates the similarity between biological replicates (Supplemental Fig. 1B). In the samples treated with EPFL6 only, we observed minimal changes in gene expression compared to the mock treatment (Supplemental Fig. 1A and B). In response to EPFL6 alone, twenty-one genes were at least two times downregulated, and ten were at least two times upregulated using a <0.05 false discovery rate (FDR) cutoff (Supplemental Table 1). Plants treated with CHX or CHX+EPFL4 clustered together but were not as similar. As expected, global inhibition of translation by cycloheximide led to widespread dysregulation of gene expression in the mock sample (Supplemental Table 1). Using the CHX treatment to control for CHX-induced effects, we analyzed the combined EPFL6 and CHX effect. Downregulated and upregulated targets of EPFL6-only treatment showed no coordinated expression pattern under CHX only treatment (Fig. 2B). Co-treatment with CHX and EPFL6 resulted in a reduction of gene expression for both downregulated and upregulated genes compared to just CHX treatment (Supplemental Table 1). This suggests that most of these genes are directly repressed by the ERf signaling pathway.

**Figure 2.**
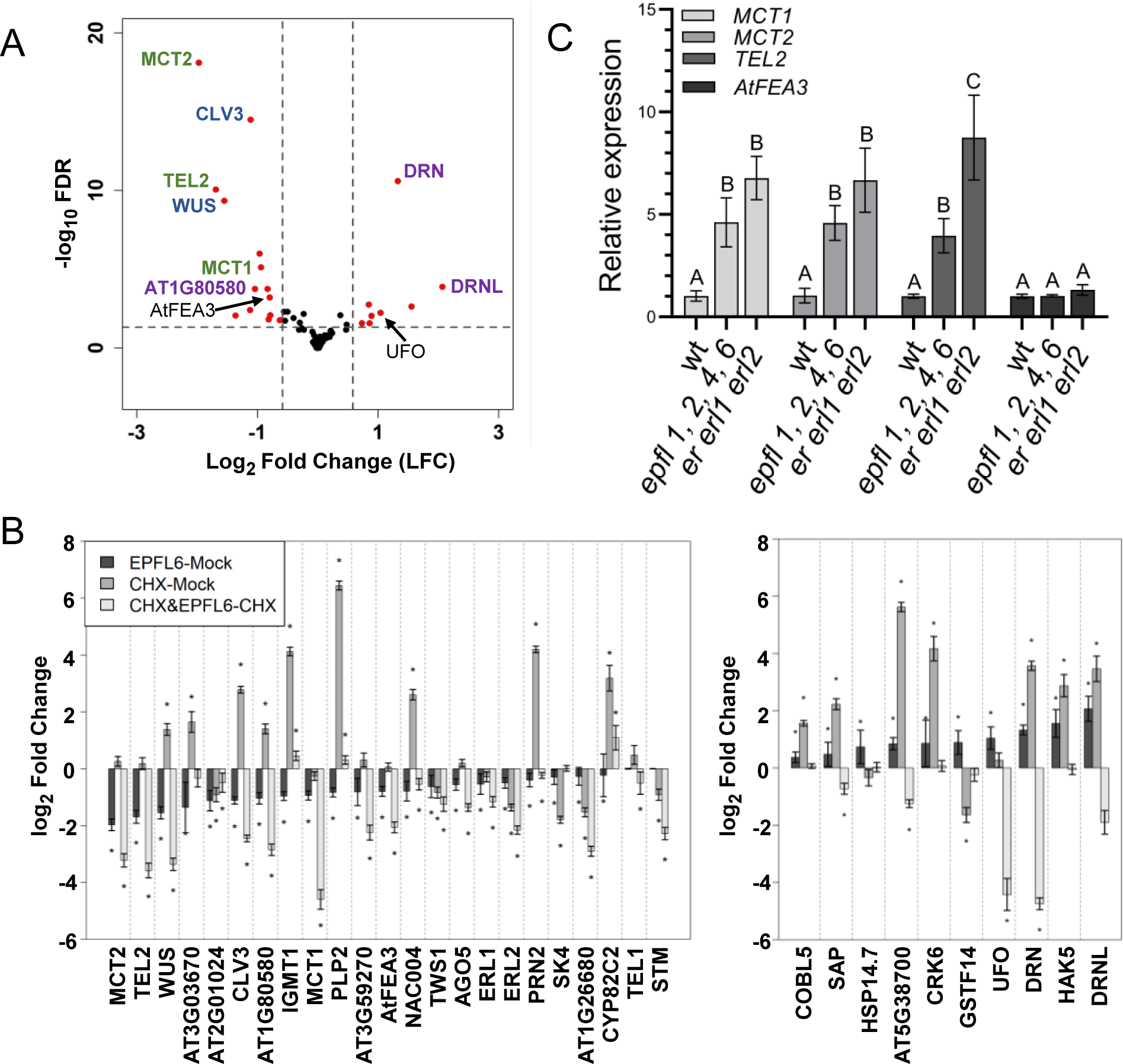
Downstream targets of EPFL6 based on RNAseq analysis. A. A volcano plot shows changes in gene expression in 3DPG *clv3 epfl1 eplf2 epfl4 eplf6* seedlings after treatment with 10 mM EPFL6. Vertical dashed lines indicate LFC cutoff =±0.585, while horizontal dashed lines the FDR cutoff =0.05. Selected genes that are discussed in the manuscript are indicated. B. Comparison of changes in gene expression in response to 10 mM EPFL6 (EPFL6-Mock), 10 mM cycloheximide (CHX-mock), and CHX and EPFL6 cotreatment versus only CHX (CHX&EPFL6-=CHX). Most genes downregulated in response to EPFL6 are also downregulated in response to EPFL6 and CHX (left panel). None of the genes upregulated in response to EPFL6 are upregulated in response to EPFL6 and CHX (right panel), suggesting that their upregulation is indirect. * FDR < 0.05, Error bars=S.E. C. Rt-qPCR analysis of selected gene expression in 3 DPG seedlings. ACTIN2 was used as an internal control. Data is average ± S.D. Statistical differences were detected using one-way ANOVA followed by Tukey post-hoc test with a minimum *P-*value of <0.05.

Consistent with our previously published data (Zhang et al., 2021), RNAseq showed that *CLV3* and *WUS* are targets of EPFL6 (Fig. 2A). Both showed ∼50% downregulation following EPFL6 treatment. The reduction in *WUS* and *CLV3* expression was independent of the production of new proteins as we observed downregulation of *WUS* and *CLV3* in samples treated simultaneously with EPFL6 and the translational inhibitor CHX (Fig. 2B). These experiments confirm that *WUS* and *CLV3* are direct targets of ERf signaling pathway in the SAM.

Additionally, we found that three members of the MEI2 family of RNA binding proteins, *MEI2 C-TERMINAL RRM ONLY LIKE 1* (*MCT1*) (AT1G37140), *MCT2* (AT5G07930), and *TERMINAL EAR-LIKE 2* (*TEL2*) (AT1G67770), were significantly downregulated by EPFL6 (Fig. 2A and B). The expression of the fourth member of this family, *TEL1* (AT3G26120), was slightly reduced in EPFL6 + CHX treatment but not by EPFL6 alone (Fig 2B). Another gene downregulated by EPFL6 was a leucine-rich-repeat protein AT3G25670 (AtFea3), one of three Arabidopsis homologs of maize *FASCIATED EAR 3* (*FEA3)* that regulates the SAM size (Je et al., 2016). RT-qPCR analysis *of er erl1 erl2* and *epfl1,2,4,6* seedings detected an increased expression of *MCT1, MCT2*, and *TEL2* which is consistent with possible ERf function in the downregulation of these genes (Fig. 2C). We were not able to consistently detect *TEL1* expression in either the wild type or the mutants due to its extremely low expression if any.

There was no change in *AtFea3* expression in the mutants (Fig. 2C), suggesting that it might not be the significant target of EPFLs. Finally, of notice is the downregulation of *ERL1* and *ERL2* expression by EPFL6. This finding agrees with previous data that ERf signaling negatively regulates *ERL1* and *ERL2* expression (Pillitteri et al., 2007).

Out of ten genes upregulated in response to EPFL6, none were upregulated when EPFL6 was applied with CHX, suggesting that upregulation of these genes is an indirect response to EPFL6. Two upregulated genes with the lowest FDR were *DORNROSCHEN (DRN)/ ENHANCER OF SHOOT REGENERATION (ESR1)* (AT1G12980) and *DRN-LIKE* (*DRNL*)/ ESR2 (AT1G24590) (Fig 2A), both APETALA2/ETHYLENE RESPONSE FACTOR (AP2/ERF) transcription factors that regulate meristem maintenance and organ initiation (Kirch et al., 2003; Ikeda et al., 2006; Ikeda et al., 2021). But both of these genes were downregulated by EPFL6 in the presence of CHX (Fig. 2B). In addition, EPFL6 downregulated AT1G80580, a close paralog of DRN and DRNL (Fig. 2A and B). This result suggests that EPFLs might downregulate this gene family directly while simultaneously indirectly promoting *DRN* and *DRNL* expression. EPFL6-only treatment upregulated the expression of *UNUSUAL FLORAL ORGANS (UFO)* (AT1G30950) (Fig. 2A), another meristematic gene (Long and Barton, 1998). However, in EPFL6+ CHX treatment, we observed the downregulation of UFO (Fig. 2 B), suggesting a complex manner in which EPFL6 regulates the expression of this gene.

EPFL2 is expressed in the boundaries between the SAM and forming primordia (Kosentka et al., 2019). To test whether EPFL2 can alter the expression of genes identified by transcriptomics from the boundary, we created *epfl1,2,4,6* transgenic plants with inducible *EPFL2* expression (*epfl1,2,4,6^T^*). We used the pOp/LhGR system that allows tissue-specific expression of a gene of choice in response to dexamethasone (DEX) (Samalova et al., 2005). The construct was created in such a way that in response to DEX, expression of EPFL2 and H2B- GFP is induced in tissues where the EPFL2 promoter is active (Supplemental Fig. 2) Before induction, we did not detect H2B-GFP. After seven hours of induction, H2B-GFP can be clearly detected in the boundary zone of the SAM (Fig. 3A). Interestingly, RT-qPCR indicated that both H2B-GFP and EPFL2 are expressed in transgenic plants without induction (Fig. 3B and C). After induction their expression increases slightly. This suggested that our construct leads to a leaky unspecific expression at low levels. This expression does not noticeably alter the phenotype of the *epfl1,2,4,6* mutant and we cannot detect H2B-GFP by microscopy. In response to DEX, GFP and presumable EPFL2 expression are strongly activated in cells where the EPFL2 promoter functions. Because this happens in very few cells out of many, RT-qPCR barely detects any change. Expression of *CLV3, MCT1*, *MCT2*, and *TEL2* genes in transgenic seedlings without induction (epfl1,2,4,6^T^-mock) was similar to their expression in untransformed *epfl1,2,4,6* (Fig. 1F, Fig. 2C, and Fig. 3D). When DEX induced EPFL2 expression for seven hours in the boundary of the SAM, expression of *CLV3, MCT1*, and *MCT2* decreased (Fig. 3D). *TEL2* might be a specific target of EPFL4 and EPFL6. Unexpectedly, very low broad expression of *EPFL2* reduced the expression of *WUS* in *epfl1,2,4,6^T^-*mock to levels that are only slightly above the wild type levels (Fig. 3E). Induction of *EPFL2* in the boundary did not significantly lower *WUS* expression. In summary, this experiment confirmed that *CLV3, MCT1*, and *MCT2* are endogenous targets of EPFL2.

**Figure 3.**
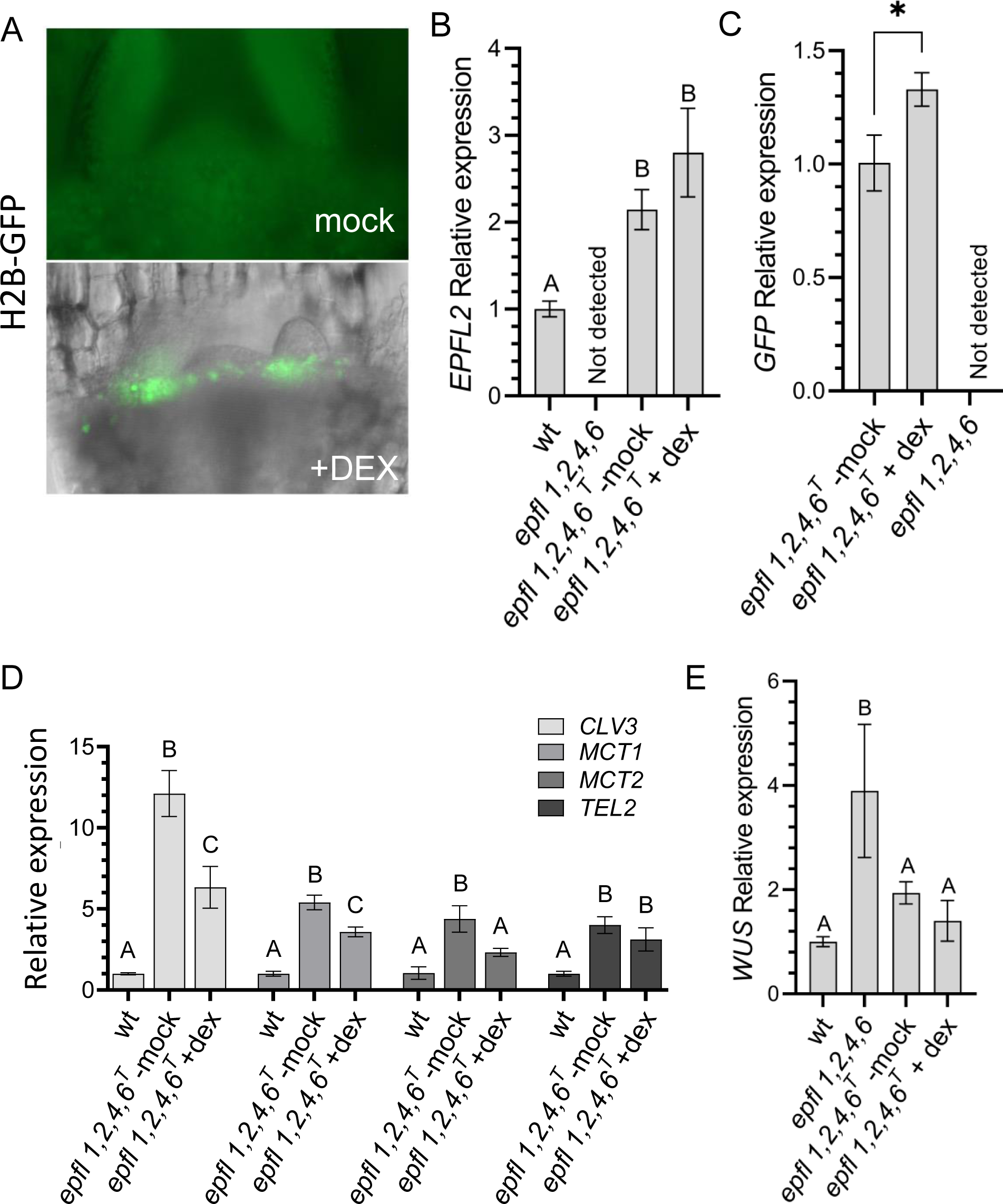
Induction of EPFL2 in the SAM boundary leads to decreased expression of *CLV3*, *MCT1*, and *MCT2*. A. Images of 3DPG *epfl 1,2,4,6* seedlings mock (an epifluorescent image) or DEX (bright filed and epifluorescent images merged) treated for 7hr. Seedlings express EPFL2 and GFP under the EPFL2 promoter that DEX induces. B-E. RT-qPCR analysis of *EPFL2, GFP, CLV3, MCT1, MCT2, TEL2,* and *WUS* expression in 3 DPG seedlings. epfl1,2,4,6^T^ are seedlings expressing inducible EPFL2. ACTIN2 was used as an internal control. Data is average ± S.D. Statistical differences were detected using one-way ANOVA followed by Tukey post-hoc test with a minimum *P-*value of <0.05.

### The expression pattern of *DRN* and *DRNL* is altered in the *epfl1,2,4,6* mutant

ERf signaling plays an important role in the initiation of cotyledons and leaves. However, the molecular mechanism is unknown (Chen et al., 2013; DeGennaro et al., 2022). Based on the role the *DRN/DRNL* gene family plays in cotyledon and leaf initiation in a variety of species (Chandler et al., 2007; Capua and Eshed, 2017; Kusnandar et al., 2021) and on our RNAseq data, we hypothesized that ERfs might regulate organogenesis through control of *DRN* and/or *DRNL* expression. To test this hypothesis, we compared the expression of their H2B-GFP reporters in the wild type and *epfl* mutants. We used a 4.9-kb sequence upstream of the start codon and a 1.4- kb sequence downstream of the stop codon to analyze *DRN* expression. These regulatory regions have been reported to reflect the *DRN* expression similarly to RNA *in situ* hybridization (Kirch et al., 2003; Luo et al., 2018). To analyze *DRNL* expression, we used a 4.3kb region upstream of the start codon as a promoter. The expression pattern of this regulatory region has also been tested previously and is consistent with RNA *in situ* hybridization (Luo et al., 2018). Based on published data, *DRN* is expressed in the young leaf primordia and the L1 and L2 layers (called the tunica) of the vegetative SAM (Kirch et al., 2003). *DRNL* is expressed in leaf and flower primordia (Ikeda et al., 2006; Nag et al., 2007). During flower development, *DRNL* is expressed in the primordia founder cells before the formation of auxin maxima and in the outer periphery of the future auxin peak (Chandler and Werr, 2014; Luo et al., 2018).

In agreement with published data, we observed *DRNp:H2B-GFP* expression in the tunica of the wild type SAM (Fig. 4A). In the L1 layer, the reporter was expressed broadly. In the L2 and deeper tissues, expression was narrow and correlated with the formation of leaf primordia.

**Figure 4.**
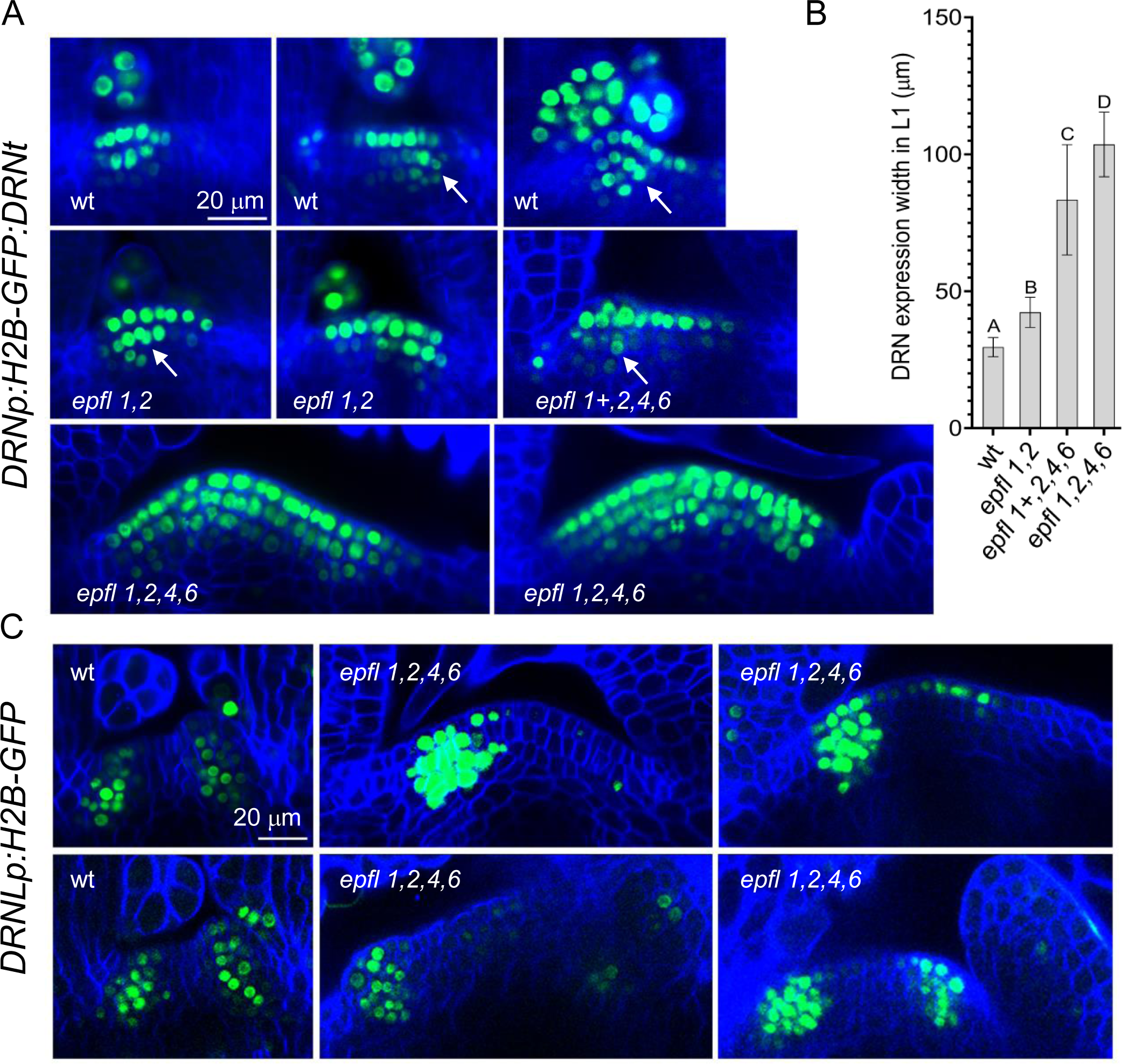
Broader expression of *DRN* and *DRNL* in the SAM of *epfl* mutants. A and C. Representative confocal images of the SAM region of 3 DPG wild-type (wt) and *epfl* mutants seedlings expressing *DRNp:H2B-GFP:DRNt or DRNLp: H2B-GFP* (green). The white arrow indicates the induction of DRN in incipient leaf primordia. In a panel, all images are under the same magnification. The cell walls were stained with SR2200 (blue). B. The average width *DRN* expression in the L1 layer of the SAM was measured on the confocal images n= 4-11. Data is average ± S.D. Statistical differences were detected using one-way ANOVA with Tukey post- hoc test with a minimum *P-*value of <0.05 and indicated by letters above bars.

We observed a similar pattern in *epfl1,2* and *epfl1/+,2,4,6* seedlings with the exception that the SAM in these mutants is slightly broader, and as a result *DRN* is expressed in a wider area of L1 (Fig 4B), and the correlation of *DRN* expression in L2 and L3 with forming organ primordia is more obvious. Even though the SAM of the *epfl1,2,4,6* mutant forms very few primordia, *DRN* is expressed very broadly in the L2 and L3 layers of this mutant. Because *DRN* is induced by auxin (Cole et al., 2009), and in the absence of ERf/EPFL signaling auxin is present at higher levels in the SAM (DeGennaro et al., 2022), it is not clear whether broader *DRN* expression in the SAM is related to altered auxin levels or if ERf signaling directly downregulates this gene.

In the wild type, *DRNL* was expressed in the narrow strip of primordia founder cells (Fig. 4C). The *epfl1,2,4,6* mutant forms leaf primordia very inefficiently (Kosentka et al., 2019). Unexpectedly, we observed efficient expression of *DRNL* in the SAM of this mutant, suggesting that in the mutant leaf primordia, founder cells are specified. This result indicates that ERf signaling promotes the subsequent step of leaf primordia outgrowth. In addition, we observed broader *DRNL* expression. While the width of the *DRNL* expression in the wild type was 2 to 3 cells, in the *epfl1,2,4,6* mutant, it was in the range of 4 to 7 cells. In the mutant, *DRNL* was also expressed in the L1 layer of the central zone. The *DRNL* promoter region contains auxin-responsive elements (Comelli et al., 2016). The broader expression of *DRNL* in the *epfl1,2,4,6* mutant can be either a direct consequence of the altered ERf signaling or due to changes in auxin accumulation.

In summary, analysis of *DRN* and *DRNL* reporters in *epfl* mutants demonstrated a broader expression of these genes. This result is consistent with the downregulation of these genes we observed under EPFL6+CHX treatment. However, this change could also result from increased auxin accumulation in the SAM of *erf* and *epfl* mutants. DRN and DRNL have been linked with the induction of leaf initiation (Chandler et al., 2007). Our data suggest that our original hypothesis that ERfs promote leaf initiation through induction of *DRN* and *DRNL* was incorrect. While increased expression of *DRN* and *DRNL* in the SAM of *epfl1,2,4,6* might alter some aspects of meristem maintenance, it is unlikely to inhibit leaf initiation. We speculate that ERfs do not specify the primordia founder cells. Their role is to promote the outgrowth of demarcated leaf primordia.

### Regulation of *WUS* expression by CLV3 signaling

In our previous work, we observed *WUS* expressed directly under the L2 layer in the wild type vegetative SAM (Zhang et al., 2021). This contradicts the accepted description of *WUS* expression in the deeper layers of the SAM (Brand et al., 2000; Schoof et al., 2000). To determine the *WUS* expression domain, we performed further analysis.

Identification of cell layers on 2D images of the SAM can be misleading because the meristem is often sectioned at an oblique angle (Supplemental Fig. 3A). During the analysis of *WUS* expression, we realized that unless we examined a 3D image, we often erroneously detected *WUS* in deeper tissue layers than it was actually expressed (Supplemental Fig. 3 B and C). Thus, we carefully analyzed z-stacks of *WUS* and *CLV3* in the wild type, *er erl1 erl2*, and *clv3*. On all images and in all seedlings, *WUS* was expressed in the third cell layer from the top at a constant distance from the surface of the SAM (Fig. 5D-G). In *clv3,* we never detected a shift of *WUS* expression upward, only a slight expansion of *WUS* downward. In the wild type, *WUS* was mostly expressed in two cell layers while in *clv3,* sometimes in three or four (Fig. 5D and E). In the *clv3 er erl1 erl2* mutant, *WUS* was primarily expressed in layers three and four from the top, with only occasional expression in deeper layers (Fig. 6). Interestingly, in this mutant *WUS* expression was discontinuous although all cells, based on CLV3 expression, are a part of the central zone (Fig. 6). In *clv3, er erl1 erl2,* and *clv3 er erl1erl2* mutants, we observed much broader expression of *WUS* along the radial axis (Fig. 5E and D, and Fig. 6) and (Zhang et al., 2021). Thus, CLV3 and ERf signaling mainly regulate *WUS* expression along the radial axis of the SAM and not the apical-basal.

**Figure 5.**
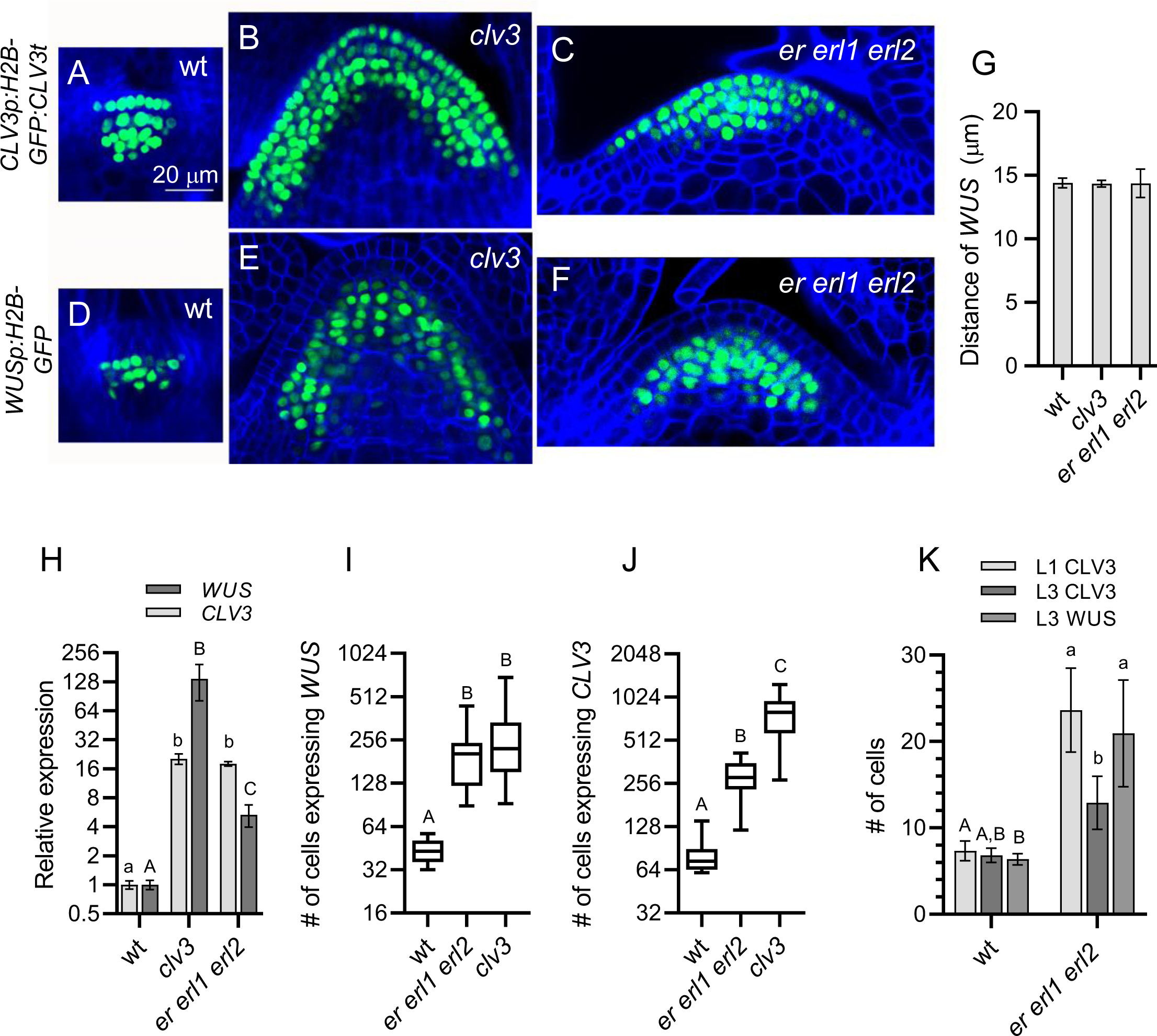
CLV3 regulates the level of *WUS* expression in the central zone but not its apical- basal pattern. A-F. Confocal images of the SAM region of 3 DPG wild-type (wt) and *clv3* seedlings transformed with *CLV3p:H2B-GFP:CLV3t* (A, B, C) and *WUSp:H2B-GFP* (D, E, F). The cell walls were stained with SR2200 (blue). All images are under the same magnification G. The distance of the *WUS* domain from the top of the SAM. *n*=4-9. H. RT-qPCR analysis of *CLV3* and *WUS* in 3 DPG seedlings of wt and mutants as indicated. *ACTIN2* was used as an internal control. I and J. The number of cells expressing *WUS* or *CLV3* in 3 DPG seedlings.n=16- 26 for *WUS* and n=12-22 for *CLV3*. K. The number of cells expressing *WUS* or *CLV3* in the L1 and L3 layer of 3 DPG SAM. n=12-21. G-K. Data is average ± S.D. Statistical differences were detected using one-way ANOVA followed by Tukey post-hoc test with a *P-*value of <0.05.

**Figure 6.**
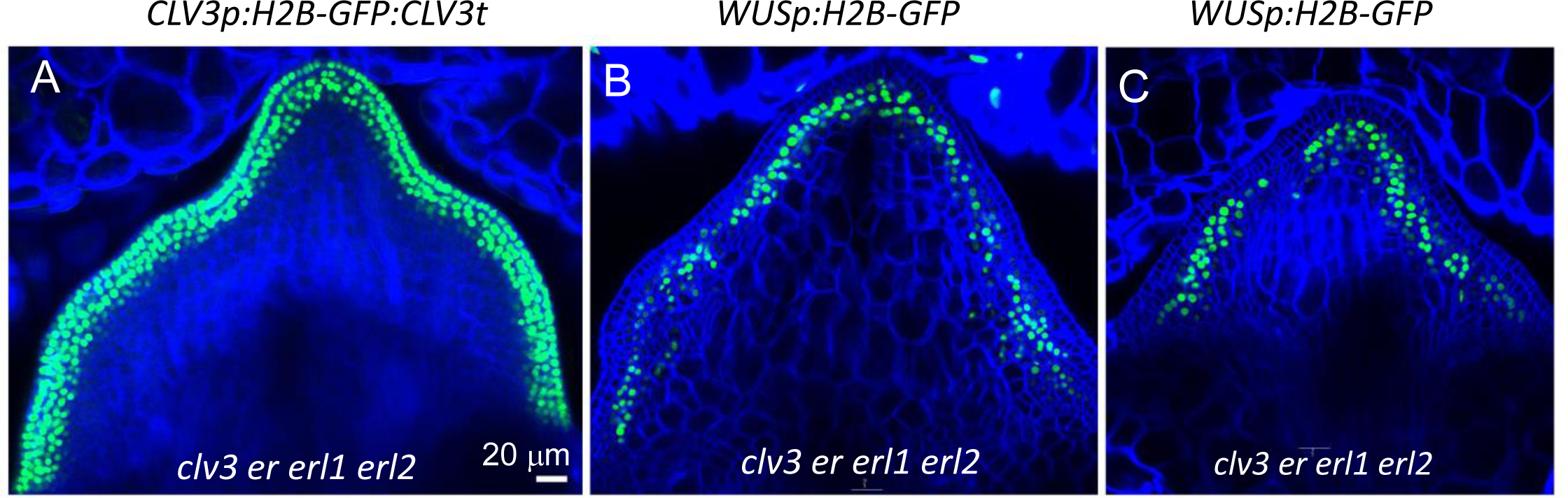
In *clv3 er erl1 erl2 mutant, c*ells that express *WUS* also express *CLV3*. A-C. Representative confocal images of the SAM region of 3 DPG *clv3 er erl1 erl2* mutant seedlings transformed with *WUSp: H2B-GFP* or *CLV3p:H2B-GFP: CLV3t* (green). In a panel, all images are under the same magnification. The cell walls were stained with SR2200 (blue).

In the vegetative SAM, *CLV3* is expressed in the top 4-5 layers, and the depth of its expression is not altered in *clv3* and *er erl1 erl2* mutants (Fig. 5A-C) (Zhang et al., 2021). Thus, in the wild type, expression of *CLV3* and *WUS* strongly overlaps. All cells that express *WUS* also express *CLV3*. This means that CLV3 should inhibit *WUS* expression primarily through autocrine signaling.

To understand how ERf and CLV3 regulate *WUS* expression, we estimated the amount of *WUS mRNA* per cell. The z-stacks were used to calculate the number of cells expressing *WUS* in the wild type and the mutants (Fig. 5I and Supplemental Videos 1-3). Compared to the wild type, there are approximately six and five times more *WUS* expressing cells in *clv3* and *er erl1 erl2*, respectively. RT-qPCR was used to determine the difference in *WUS* expression (Fig. 5H). After the difference in the number of *WUS* positive cells was taken into account, the RT-qPCR indicated that individual cells in *clv3* expressed approximately 22 times more *WUS*. In contrast, the amount of *WUS* per cell is not significantly changed in *er erl1 erl2* (Table 1). This result indicates that the function of CLV3 is to regulate the levels of *WUS* expression in cells of the central zone. On the other hand, ERfs restrict the *WUS* expression domain in the SAM periphery but do not control the cellular levels of *WUS* in the central zone. In addition, we observed a considerable variance in the number of *WUS* positive cells in individual meristems of both mutants (Fig. 5I), suggesting that both CLV3 and ERfs are necessary for the SAM size stability.

**Table 1.** Comparison of *CLV3* and *WUS* expression in the wild-type (WT), *clv3*, and *er erl1 erl2* 3DPG seedlings. The number of cells was determined on composite images obtained by confocal microscopy. The changes in gene expression per cell in mutants were determined by dividing the fold increase in gene expression as determined by RT-qPCR by the fold change of the number of nuclei expressing the gene.

| | # of cells with<br>*WUS* | ≈ change in<br>*WUS/cell* | # of cells with<br>*CLV3* | ≈ change in<br>*CLV3/cell* | Ratio of *CLV3*<br>cells/ *WUS* cells |
| --- | --- | --- | --- | --- | --- |
| WT | 43.5 ± 8.0 | 1x | 81.7±23.9 | 1X | 1.9X |
| *clv3* | 265.8 ± 151.3 | 22x | 791.7 ± 258.3 | 2.1X | 3.0X |
| *er erl1 erl2* | 208.8 ± 97.0 | 1.1x | 286.4 ± 73.7 | 5.1X | 1.4X |

*Clv3-9* has a point mutation that results in a premature stop codon (W62STOP), but it can still produce *CLV3* mRNA. Compared to the wild type, there are approximately 9.7 and 3.5 times more *CLV3* expressing cells in *clv3* and *er erl1 erl2*, respectively (Fig. 5J). Surprisingly, a dramatic ∼22x increase in *WUS* in *clv3* leads only to a relatively modest ∼2.1x increase in *CLV3* (Table 1). One possibility is that a premature stop codon decreases *CLV3* mRNA stability. On the other hand, it was previously proposed that at high concentrations, WUS can inhibit *CLV3* expression (Perales et al., 2016). While individual meristematic cells in *er erl1 erl2* do not have increased *WUS* expression, they accumulate approximately five times more *CLV3* (Table 1), suggesting that ERfs regulate *CLV3* expression in individual cells independently of *WUS*.

Next, we investigated whether in *clv3* and *er erl1 erl2* mutants there was a comparable increase in the number of cells expressing *WUS* and *CLV3*. In the wild type, there are 1.9 times more cells expressing *CLV3* than cells expressing *WUS* (Table 1). This is consistent with the fact that *CLV3* is expressed in almost all *WUS*-expressing cells plus tunica cells. In the *clv3* mutant, the ratio of *CLV3* cells to *WUS* cells increases from 1.9x to 3x due to faster cell proliferation of the tunica cells. This leads to the convex shape of the *clv3* SAM. We were surprised that in the *er erl1 erl2* mutant the ratio of *CLV3* expressing cells to *WUS* expressing cells decreased from 1.9x to 1.4x. To understand the cause of this decrease, the expression of both genes was compared in individual cell layers of the wild type and *er erl1 erl2* (Fig. 5K). In the mutant, *CLV3* expanded very broadly in the L1 layer. However, it did not spread as widely in the internal tissues as *WUS*. This finding has several implications. First, it suggests a complex tissue-specific pattern in which *CLV3 e*xpression is regulated and suggests that ERf signaling plays an especially strong role in the inhibition of *CLV3* expression in the L1. An additional mechanism might restrict *CLV3* expression in the internal tissues of the meristem periphery. Second, uneven expansion of *CLV3* and *WUS* domains in the *er erl1erl2* mutant indicates that the changes in their expression are not due to the overall expansion of the central zone but are instead due to a particular mechanism by which ERfs regulate them.

### A mechanistic model for 3D expression patterning in the SAM

Previous mathematical models for apical-basal patterning of gene expression in the SAM either assumed or produced antiparallel gradients of *WUS* and *CLV3* expressions with minimal overlaps (Hohm et al., 2010; Chickarmane et al., 2012; Liu et al., 2020), which contradict our 3D high-resolution imaging data (Fig. 5A-F and Supplemental Videos 1-3). To test whether our current understanding of the regulatory network involving WUS, CLV3, and ERf signaling is sufficient to explain the up-to-date expression data, we built a 3D mathematical model that describes both the steady state geometry of the SAM in terms of cell locations and the expression regulations of *WUS* and *CLV3*. In this reaction-diffusion model, cells in the SAM were represented as 326 points in a 3D half dome. For gene regulations that occur in cells, we considered the canonical negative feedback loop between WUS and CLV3 (Brand et al., 2000), the negative regulation of both *WUS* and *CLV3* by EPFL whose expression zones were restricted to the peripheral areas of the SAM (Kosentka et al., 2019; Zhang et al., 2021) and the negative regulation of *CLV3* by HAM signal from the meristem rib (Fig. 7A) (Han et al., 2020b). For *WUS* and *CLV3*, mRNA and protein levels were modeled separately. The model also considered movements of WUS, CLV3, and EPFL between neighboring cells. Finally, we considered the inhibition of *CLV3* by high concentrations of WUS (Perales et al., 2016) (Fig. 7A. Dash line).

**Figure 7.**
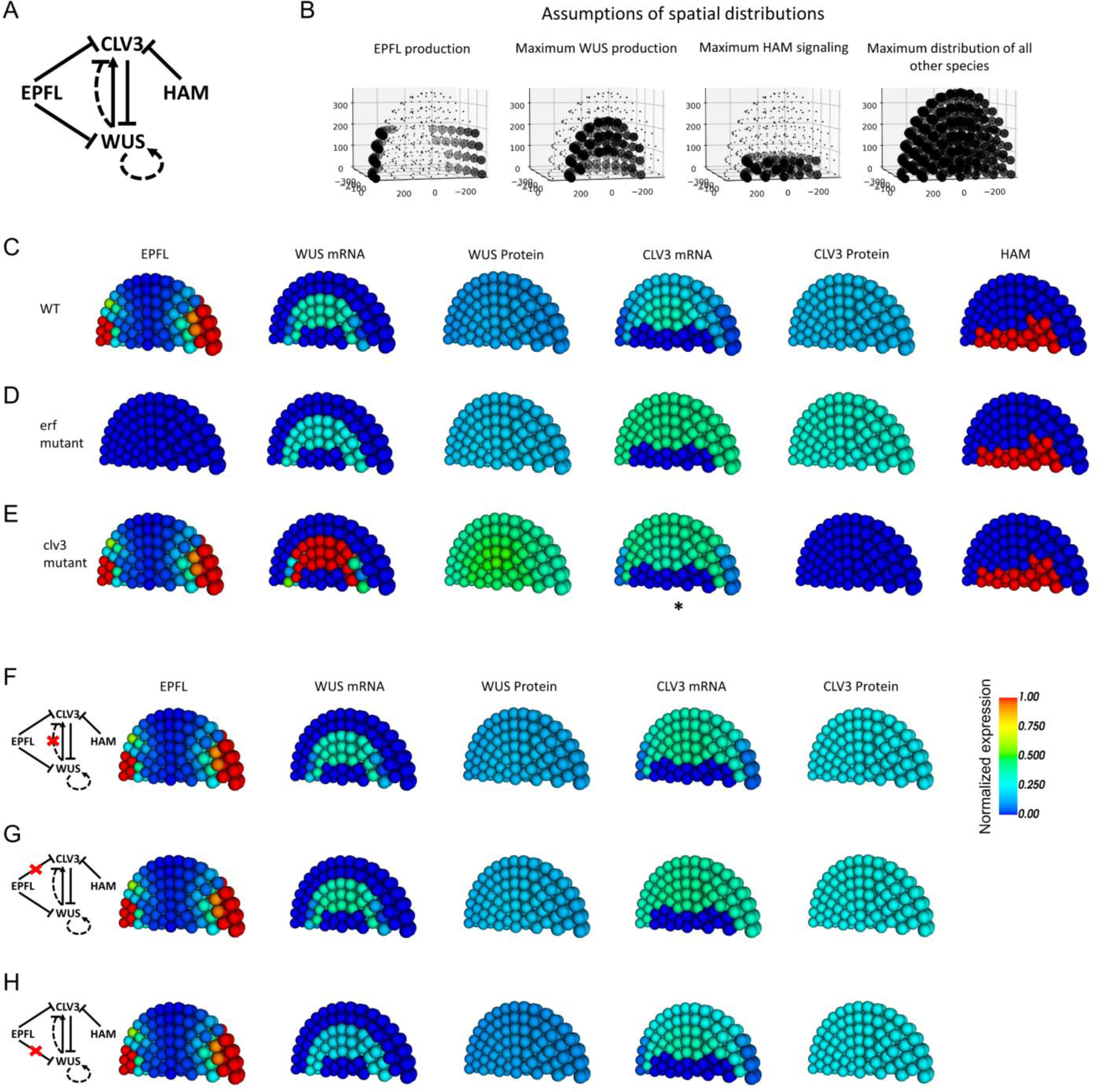
A mathematical model of SAM patterning. A. General gene regulatory network describing transcriptional regulations in the SAM. For *CLV3* and *WUS*, both mRNA and protein are explicitly described, whereas only proteins were explicitly described for EPFL and HAM. B. Assumptions on spatial distributions of regulatory factors. Dots show the positions of simulated cells. Small dots mean the absence of the factor from the regulatory network in A. C-E. Model simulations under experimental conditions are included in this study. The colors of the balls show normalized expression levels of indicated factors (see Methods). *: *CLV3* mRNA is nonfunctional and not translated. F-H. Additional model conditions for predictions on the roles of specific transcriptional regulations.

We restricted the *WUS* expression to L3 and lower layers and the HAM signal to the 6^th^ layer from the epidermis and below to account for other spatial factors (e.g. cytokinin receptor) not described in the model (Fig. 7B).

We fit the model to our high-resolution experimental data using biologically plausible parameter values. The model reproduced the *CLV3* expression region encompassing the *WUS* expression region under the wild type condition (Fig. 7C). This substantial overlap was also observed with the *erf* mutant (the removal of EPFL signal) and the *clv3* mutant (note that in the latter case the *CLV3* mRNA is produced but nonfunctional). Compared to previous models, our model does not require an assumption of a HAM signal in layers L1-L5 close to the epidermis in the SAM (Figure 7C-E, last column) (Zhou et al., 2018; Liu et al., 2020). In addition to the *CLV3*-*WUS* overlap, the absence of EPFL signal resulted in an expansion of the *WUS* expression region, but the level of expression in *WUS* mRNA-containing cells was unchanged (Fig. 7D). In contrast, the absence of functional CLV3 gave rise to both expansion of expression region and single-cell upregulation of *WUS* (Fig. 7E). The expression of *CLV3* in both mutants was expanded in the SAM, but its single-cell upregulations were much less prominent compared to *WUS* (Fig. 7D and E).

We found that the loss of *CLV3* inhibition by high concentration of WUS resulted in both expansion and single-cell upregulation of CLV3 expression (compare Fig. 7F and C). We next asked whether the inhibitions of both *CLV3* and *WUS* by EPFL are required for correct SAM patterning. The removal of *CLV3* inhibition by EPFL resulted in upregulation of *CLV3* in single cells and expansion of *CLV3* expression (Fig. 7G), whereas the removal of *WUS* inhibition by EPFL resulted in expansions of expression regions of both *CLV3* and *WUS* (Fig. 7H). This suggests that the two regulations by EPFL are required for SAM patterning. Taken together, our experimentally inspired 3D model reproduced key quantifications and distributions of gene expressions with fewer assumptions compared to previous models, and it suggests new mutant phenotypes that can be tested in future experiments.

## DISCUSSION

### The role of ERf/EPFL signaling in the regulation of *CLV3* and *WUS* expression

ERf receptors are important negative regulators of SAM size, functioning through suppression of *CLV3* and *WUS* expression (Uchida et al., 2012a; Chen et al., 2013; Uchida et al., 2013; Mandel et al., 2014; Zhang et al., 2021). The analysis of transcriptome changes after brief activation of ERf signaling confirmed that these two genes are the main targets of the pathway. A mechanistic model for 3D expression patterning in the SAM indicated that regulation of both genes is necessary for the correct SAM patterning. The three ERf receptors redundantly control the vegetative SAM’s width and are particularly important during embryogenesis when the meristematic domain is defined (Uchida et al., 2012a; Chen et al., 2013). This is in contrast to CLV3 signaling, which controls both the width and the height of the SAM and functions in maintaining the meristem at a relatively constant size throughout a plant’s life (Clark et al., 1995). Consistent with their different roles in SAM establishment and maintenance, ERfs and CLV3 play distinct roles in the regulation of *WUS*. ERfs regulate the width of the *WUS* domain, suppressing its expression in the periphery of the SAM. In other words, ERf signaling contributes to the patterning of the SAM, defining different zones. In contrast, CLV3 regulates the cellular concentration of *WUS* in cells of the central zone, which defines the size of the central zone.

The second function of ERfs is to inhibit *CLV3* expression. ERfs suppress CLV3 cellular levels and prevent its expression in the SAM’s periphery. This suppression is especially strong in the L1 layers of the SAM. ERfs do this independently of *WUS*, since in the central zone of the *er erl1 erl2* mutant the *CLV3* cellular expression increases without an increase of *WUS* expression, and knockout of ERf signaling in the *wus* background promotes *CLV3* expression (Kimura et al., 2018). In meristematic cells of bryophytes, CLV signaling regulates auxin and cytokinin signaling, and it does it independently of *WUS* homeobox-containing (*WOX*) genes (Fouracre and Harrison, 2022). CLV3 signaling might have additional unknown targets in angiosperms, and suppression of CLV3 by ERfs might be related to these other functions.

The activity of ERf receptors is regulated by small extracellular cysteine-rich proteins from the EPF/EPFL family. In Arabidopsis, this family consists of eleven genes that form four clades (Takata et al., 2013). The function of two clades is linked with the formation of stomata (Richardson and Torii, 2013). The other two clades regulate the SAM (Kosentka et al., 2019). There are differences in these two last clades’ expression and overall function. One clade, consisting of EPFL4, EPFL5, and EPFL6, promotes elongation of aboveground organs. EPFL4 and EPFL6 are expressed in the endodermis and regulate the elongation of internodes and pedicels (Abrash et al., 2011; Uchida et al., 2012b). All three genes redundantly promote the elongation of stamen filaments (Huang et al., 2014; He et al., 2023; Negoro et al., 2023).

Another clade consists of EPFL1, EPFL2, and EPFL3. EPFL2 regulates ovule initiation, elongation of leaf teeth, and growth of cotyledons, and it is often expressed in organ boundaries (Tameshige et al., 2016; Kawamoto et al., 2020; Fujihara et al., 2021). EPFL4/5/6 and EPFL1/2/3 differ not only in function but also in structure. All EPF/EPFLs are composed of a loop and a scaffold (Ohki et al., 2011). The loop structure is important for ligand function and might define whether it is an agonist or antagonist. EPFL9 is an antagonist; it competes with EPF2 for the same receptors but does not activate the downstream MAPK cascade (Lee et al., 2015). Swapping the loops between EPF2 and EPFL9 reverses their function (Ohki et al., 2011). The sequence and length of loops differ significantly between EPFL1/2/3 and EPFL4/5/6 clades. EPFL1, EPFL2, EPFL4, and EPFL6 redundantly regulate the SAM width, leaf initiation, and internode elongation (Kosentka et al., 2019). But do they regulate the same set of genes?

Previously, we demonstrated that EPFL4 and EPFL6 inhibit the expression of *WUS* and *CLV3* (Zhang et al., 2021). Our current work shows that EPFL1 and EPFL2 also regulate the expression of these two genes. In summary, while the two clades of EPFL ligands differ structurally, their function in the SAM is very similar.

Previously, we observed that while the SAM of *clv3 er erl1 erl2* forms very few organs, the SAM of *clv3 epfl1 epfl2 epfl4 epfl6* forms some leaves and flowers (Zhang et al., 2021). A current comparison of *WUS* and *CLV3* expression in *er erl1 erl2* and *epfl1 epdl2 epdl4 epfl6* detected some small but statically significant differences. This finding suggests that either ERf receptors regulate the SAM in a ligand-independent manner or other EPFLs also regulate the SAM structure.

### ERf/EPFL signaling does not designate cells for leaf primordia but promotes primordia outgrowth

ERf signaling promotes the initiation of cotyledons and leaves and regulates phyllotaxis (Chen et al., 2013; DeGennaro et al., 2022). The hormone auxin induces initiation of aboveground organs. However, in the absence of ERf signaling, auxin cannot initiate leaves and cotyledons efficiently (DeGennaro et al., 2022). Previously, we proposed that auxin and ERfs have common downstream targets (DeGennaro et al., 2022). In Arabidopsis, two AP2 transcription factors, *DRN* and *DRNL,* promote cotyledon and leaf initiation (Chandler et al., 2007). During organogenesis, *DRNL* is expressed in incipient organ primordia before the formation of auxin response maxima, and it functions synergistically with auxin and PID (Chandler et al., 2011). DRNL in complex with transcription factor MONOPTEROS (MP) inhibits cytokinin accumulation in forming primordia (Dai et al., 2023). RNAseq data indicated that in the absence of CHX, a brief activation of ERf signaling promotes *DRN* and *DRNL* expression; however, in the presence of CHX, it downregulates *DRN* and *DRNL* expression. To understand the role of ERf signaling in the regulation of these two genes, we analyzed their expression in the SAM. This analysis indicated that *DRN* and *DRNL* are expressed more broadly in the *epfl 1,2,4,6* mutant than in the wild type, which is consistent with the downregulation of *DRN* and *DRNL* by EPFL6 in the presence of CHX. Overall, these data indicate that cells designated to become leaf primordia are specified, but some other requirements for leaf primordia outgrowth still must be met.

Transcriptomic analysis identified a family of mei-2-like RNA-binding proteins as additional putative targets. Nine mei-2-like genes in Arabidopsis form two clades (Anderson et al., 2004). EPFL6 inhibits the expression of three genes belonging to the same clade: *MCT1*, *MCT2*, and *TEL2*. Based on *in situ* hybridization, all three of these genes are expressed in the central zone of the SAM (Anderson et al., 2004; Yadav et al., 2009). In maize, rice, and moss, these genes inhibit leaf initiation and control phyllotaxy (Veit et al., 1998; Kawakatsu et al., 2006; Xiong et al., 2006; Vivancos et al., 2012), but their meristematic function in Arabidopsis is unknown. It is tempting to speculate that ERfs regulate organogenesis by inhibiting *MCT/TEL* family gene expression. However, analysis of these genes’ function in Arabidopsis is necessary before any definitive conclusion can be made.

### CLV3 controls *WUS* cellular levels but not the apical-basal position of its expression domain

In Arabidopsis, the WUS-CLV3 negative feedback loop is central to the stability of the SAM size (Han et al., 2020a). WUS promotes the identity and proliferation of stem cells. CLV3 inhibits the expression of *WUS* to prevent their overproliferation. Because *WUS* positively regulates *CLV3* expression, CLV3 signaling can decrease *WUS* expression only to a certain extent and can never completely shut it down. The prevailing model asserts that the function of CLV3 is not only to regulate cellular levels of WUS but also to position the *WUS* expression domain in the deeper tissues of the SAM. This model was proposed during early investigations of the CLV3 and WUS feedback loop. The comparison of *WUS* expression by *in situ* hybridization in the wild type and in the *clv3* mutant was interpreted as *WUS* being expressed in deep layers of the SAM, but when *CLV3* is absent, its expression moves upward directly under the L2 cell layer (Brand et al., 2000; Schoof et al., 2000). However, multiple published *in situ* hybridization images show that in the wild type, *WUS* is expressed directly under the L2 layer in both the vegetative and inflorescence SAM (Mayer et al., 1998; Lenhard and Laux, 2003; Hu et al., 2018; Luo et al., 2018). In addition, the original manuscript by Schoof and colleagues states that in embryos, *WUS* is expressed directly under the L2 layer in the wild type, and its expression domain does not move upward in the *clv3* mutant (Schoof et al., 2000). An examination of *in situ* images of *WUS* expression in the manuscript by Brand and colleagues shows *WUS* expression directly under the L2 layer in the wild type (Brand et al., 2000). In wild type meristems, the presence of *WUS* mRNA directly under the L2 layer was observed for *pWUS:GFP:WUS*, *pWUS:WUSlinker-GFP,* and *pWUS:2xVenus-NLS:tWUS* constructs (Yadav et al., 2011; Daum et al., 2014; Wenzl and Lohmann, 2023). Despite all this evidence, the assumption that CLV3 regulates *WUS* expression along the apical-basal axis has not been explicitly challenged.

The correct detection of *WUS* expression on *in situ* hybridization images depends on the precise vertical sectioning of the SAM exactly through the center of the SAM. If the section is made at an oblique angle, the *WUS* expression will be perceived to be deeper than it is. We analyzed *WUS* expression using the *WUSp:H2B-GFP* construct. This construct contains a 4.5 kb regulatory sequence that has been used previously and that includes all regulatory elements necessary for expression in the SAM (Bäurle and Laux, 2005; Yadav et al., 2009; Zhang et al., 2017). When we examined two-dimensional images, we realized that they provide an inconsistent pattern of *WUS* expression and are challenging to interpret. However, analysis of z- stacks firmly placed *WUS* expression in the third and fourth cell layer of the SAM in the wild type and in any mutant that we observed. There was no shift of *WUS* expression apically in *clv3* or other mutants. If there was any expansion of *WUS* expression along the apical-basal axis, it was basally into the fifth layer in some *clv3* and *epfl* seedlings. The expression of *WUS* is induced by cytokinins, which are produced in the L1 layer of the SAM but are perceived only underneath the tunica (Lindsay et al., 2006; Gordon et al., 2009; Chickarmane et al., 2012).

Diffusion of cytokinins from the epidermis tethers *WUS* expression to a specific distance from the apex; there is no apparent need for additional regulation. Our data indicates that CLV3 does not define the *WUS* expression domain but controls the concentration of *WUS* in the central zone.

Based on published *in situ* hybridization images, expression of *CLV3* varies and can be detected in either three or four top layers of the SAM (Fletcher et al., 1999; Brand et al., 2000; Lenhard and Laux, 2003; Reddy and Meyerowitz, 2005). This inconsistency could be due to differences in the SAM sectioning (through the middle or at an angle), differences between vegetative, inflorescence, and floral meristems, or variable growth conditions. Recent findings suggest that the depth of *CLV3* expression is regulated by temperature, with higher temperatures inhibiting *CLV3* expression in the deeper tissues (Wenzl and Lohmann, 2023). Our data indicate that in the vegetative SAM at 21°C, *CLV3* is expressed in four top cell layers in the wild type, *clv3*, *er erl1 erl2*, *epfl,* and *clv3 er erl1 erl2* mutants, and its expression strongly overlaps with *WUS*. Thus, in the vegetative SAM, *CLV3* should strongly regulate *WUS* cellular levels through autocrine signaling, which is typical for CLAVATA3/EMBRYO SURROUNDING REGION-RELATED (CLE) peptides (Narasimhan and Simon, 2022). Modeling predicted that the overlap of *CLV3* and *WUS* expression removes the necessity for HAM influence close to the epidermis in the SAM. We observed that *CLV3* expression in L1 and deeper tissues is controlled differently.

CLV3 may be suppressed in the deeper tissues of the SAM during bolting or in response to changing environmental conditions, and that is why it was not detected there in some experiments. Further research on the mechanisms controlling *CLV3* expression, including the role of autocrine signaling versus signaling from the tunica, should provide deeper insights into the molecular processes that oversee the size of the SAM and impact the overall plant architecture.

## MATERIALS AND METHODS

### Plant Materials and Growth Conditions

The Arabidopsis thaliana ecotype Columbia was used as the wild type. The following mutants have been described previously: *er-105 erl1-2 erl2-1* (Shpak et al., 2004), *epfl1-1 epfl2- 1 (*abbreviated here as *epfl1,2), cll2-1 chal-2/epfl4 epfl6 (*abbreviated here as *epfl4,6), epfl2,4,6,* and *epfl1,2,4,6* (Kosentka et al., 2019), *clv3-9* (Nimchuk et al., 2015), and *clv3 epfl1,2,4,6* (Zhang et al., 2021). All mutants are in the Columbia background. Seedlings were grown on modified Murashige and Skoog medium plates supplemented with 1% (w/v) sucrose. Plates were stratified for two days at 4°C and then moved to a growth room with the following conditions: 18 h light/6 h dark cycle and 21°C. The generation of the wild type plants expressing *WUSp:H2B- GFP:35St* (pESH746) and *CLV3p:H2B-GFP:CLV3t* (pESH 747) constructs was described previously (Zhang et al., 2021). These constructs were transformed into *clv3, epfl1,2, epfl4,6*, *epfl1/+,2,4,6,* and *clv3 er erl1 erl2* plants.

The DRNp:H2B-eGFP:DRNterm construct was generated by fusing the 4.86 kb sequence upstream of the start codon with H2B-eGFP followed by a 1.38 kb sequence downstream of the stop codon (Kirch et al., 2003; Luo et al., 2018). H2B-eGFP was fused to the downstream sequence using overlapping PCR and inserted into the binary vector pPZP222 between BamHI and SalI sites. The template for amplifying the H2B-EGFP sequence was a plasmid from the Z. Nimchuk lab (UNC Chapel Hill, USA). The H2B-eGFP:DRNterm plasmid was used as a vector to introduce the 4.86 kb promoter sequence using BamHI, and the plasmid was named pAMO102b. The DRNLp: H2B-eGFP:35sT construct was generated by fusing the 4.3 kb sequence upstream of the start codon to H2B-eGFP:35sTerm (Luo et al., 2018). Using overlapping PCR, H2B-eGFP was linked to the CaMV 35s terminator and inserted into the binary vector pPZP222 between BamHI and SalI sites. This plasmid was used as a vector to introduce the 4.3 kb DRNL promoter between KpnI and BamHI sites, and the construct was named pAMP109. Both constructs were confirmed by Sanger sequencing.

The following construct was generated to produce dexamethasone-inducible EPFL2 expressed in its endogenous domain (Supplemental Fig. 2). The 2.58 kb promoter of EPFL2 was amplified using pPZK412 (Kosentka et al., 2019) and fused with GR-LhG4:T35S amplified from pBIN-LhGR-N (Samalova et al., 2005) via overlapping PCR. This DNA fragment was introduced into the binary vector pPZP222 between KpnI and SbfI sites, and the plasmid was named pAMO112. The next step involved 8 PCR reactions. PCR1 amplified H2B-GFP:35S terminator (T_35S_). PCR2 amplified the omega translational enhancer (Ω) and 35s minimal promoter (35S Min) using plasmid pH-TOP (Samalova et al., 2005) as a template. PCR1 and PCR2 products were overlapped to generate a PCR3 fragment. PCR4 used pH-TOP as a template to generate a fragment containing pPOP6 followed by 35S Min and Ω. PCR5 amplified the EPFL2 coding sequence with introns using the pPZK412 vector as a template. PCR6 amplified the NOS terminator (T_NOS_) using pBIN-LhGR-N as a template. PCR7 was an overlapping PCR that fused DNA fragments created by PCR4, PCR5, and PCR6. Finally, PCR8 was an overlapping PCR that fused the PCR3 and PCR7 DNA fragments. It generated a DNA fragment containing T_35s_:H2B-GFP: Ω:35SMin:pOp6:35SMin: Ω:gEPFL2: T_NOS_. The DNA segment generated by PCR8 was inserted into the SbfI site of pAMO112. Orientation was confirmed via restriction digest and Sanger sequencing. The sequencing identified additional SbfI sites between two T35s. The final construct was named pAMO113 and contained: proEPFL2:GR- LhG4:T_35S_ T_35S_:H2B-GFP:Ω:35sMin_pOp6_35sMin:Ω:gEPFL2:T_NOS_.

The generated constructs were transformed into an Agrobacterium tumefaciens strain GV3101/pMP90 by electroporation and introduced into the wild type (Columbia ecotype), *epfl1,2* and *epfl1/+,2,4,6* plants by the floral dip method.

To induce EPFL2, an *epfl1,2,4,6* mutant expressing pAMO113 was grown on modified Murashige and Skoog medium (MS-0) plates for five days (3DPD). Then 15 seedlings per biological replicate were transferred to 2 ml of liquid MS-0 containing 10 µM Dex or an equivalent amount of DMSO (mock treatment). The 50 mM stock solution of Dex was prepared using DMSO. 24 well-cultured plates with the samples were kept on a rocker in a growth room for seven hours, and then seedlings were preserved in liquid nitrogen.

Due to the infertility of the *epfl1 epfl2 epfl4 epfl6* mutant, we isolated it from the progeny of *epfl1/+ epfl2 epfl4 epfl6* plants. A small piece of root was cut from a seedling and placed into the PCR mix. PCR was performed using the Phire Plant Direct PCR Master Mix kit (Thermo Fisher Scientific). The rest of the seedling was preserved in 4% paraformaldehyde for microscopy. A three-primer PCR reaction with EPFL1.74, EPFL1.436.rev, and 3dspm was performed to genotype for *epfl1*. The mutant band was approximately 200bp, and the wild type band 387bp.

### RNA-Seq library construction, sequencing, and analysis of differential gene expression

For the RNA-Seq sample collection, seedlings were grown as described previously (Zhang et al., 2021). In brief, the *clv3 epfl1,2,4,6* seedlings were grown on modified MS medium plates for five days (3DPG). Four treatment conditions were used: mock, 10μM EPFL6, 10μM cycloheximide, and a combination of 10μM EPFL6 and 10μM cycloheximide. Purification of the EPFL6 peptide was described previously (Lin et al., 2017). EPLF6 peptides were diluted in 10mM Bis-Tris, 100nM NaCl, pH-6.0 (treatment buffer). For treatment, 10 seedlings per biological replicate were transferred into 1 ml of liquid MS medium. Seedlings in liquid medium were treated with 8.7μl of the 1.15 mM EPFL6 solution or for mock with an equal volume of the treatment buffer. For cycloheximide treatments, seedlings in a liquid medium were pretreated with 10μM cycloheximide for 10 minutes before EPFL6 or mock was added. The aboveground portion of the seedlings was collected 3 hours after adding EPFL6 or the mock treatment buffer and flash frozen in liquid nitrogen. Three biological replicates were performed for each treatment. Total RNA was isolated using the Spectrum Plant RNA Isolation Kit (Sigma-Aldrich).

RNA quality was measured using a bioanalyzer; all samples had a RIN score greater than 7.5. Paired-end cDNA libraries were constructed using the TruSeq mRNA kit from Illumina.

The libraries were sequenced on a NovaSeq S4 flow cell in paired-end mode and with 150 base pair long reads at the Oklahoma Medical Research Foundation. Raw read quality was assessed with FastQC v0.11.5. Raw reads were aligned to the TAIR10.1 genome and Araport11 annotation (TAIR genome and Araport11 citation) using STAR-2.7.6a (Dobin et al., 2013), with default parameters except for the following: - alignIntronMax 1000. Mapping quality was assessed with RSeQC v4.0.0 (Wang et al., 2012). Reads were counted using subread featureCounts v2.0.1 (Liao et al., 2013) in paired-end mode. Reads were imported into R (v3.6.3). Genes not expressed in all three replicates of at least one sample were removed.

Samples were inspected for batch effect by PCA, and no batch effect was found. The filtered reads were then normalized, and differential gene expression was assessed using DESeq2 v1.26.0 (Love et al., 2014) using a two-factorial design. The resulting p-values were corrected for multiple comparisons using FDR, and the resulting log2 fold changes were shrunk using ashr2.2 (Stephens, 2017).

### Quantitative RT-PCR analysis

Total RNA was isolated from the tissues of 3DPG seedlings using the Spectrum Plant RNA Isolation Kit (Sigma-Aldrich). The RNA was treated with RNase-free RQ1 DNase (Promega). First-strand complementary cDNA was synthesized with LunaScript™ RT SuperMix Kits (New England Biolabs). Quantitative PCR was performed with a CFX96 Touch Real-Time PCR Detection System (Bio-Rad) using SsoAdvanced Universal SYBR Green Supermix (Bio-Rad). Each experiment contained three technical replicates of three biological replicates. Cycling conditions were as follows: 30s at 95°C; then 40 repeats of 10s at 95°C, 10s at 52°C for *ACTIN2*, 55°C for *WUS* and *GFP*, 53°C for *CLV3*, 56.7°C for *MCT1*, 50°C for *MCT2* and *EPFL2*, and

56.1°C for TEL2, and 15s at 68°C, followed by the melt-curve analysis. For AtFEA3, two-step PCR was performed. Cycling conditions were as follows: 30s at 95°C, then 40 repeats of 10s at 95°C, 30s at 68°C, followed by the melt-curve analysis. qPCR for ACTIN, CLV3, EPFL2, and GFP was performed in 10 μl with 4 µl of 10× diluted cDNA, while WUS, AtFEA3, MCT1, MCT2, and TEL2 were performed in 20 μl with 8 µl of 10× diluted cDNA reaction. All primers used in this study are in Supplemental Table 2. The fold difference in gene expression was calculated using relative quantification by the 2−ΔΔCT algorithm.

### Microscopy

For microscopy, we used the T3 or T4 generations of transgenic plants that were homozygous for the insert, except when the *WUS* reporter was analyzed in T2 *epfl 1,2* and *epfl 4,6* seedlings. Three-day-old seedlings were fixed with 4% paraformaldehyde for 1.5h. The fixed samples were washed three times for 5 min in phosphate buffer (PBS) and cleared with ClearSee (Kurihara et al., 2015) for three days at room temperature on a rocker. The cell wall was stained with Renaissance 2200 [0.1% (v/v) in ClearSee] (Musielak et al., 2015) for 1–2 days. For better imaging, one cotyledon was removed under a stereo microscope. A Leica SP8 Confocal microscope (Leica Microsystems, Wetzlar, Germany) was used at the UTK Advanced Microscopy and Imaging Center. An argon laser with 488 nm emission was used for the excitation of EGFP, and images were collected using a HyD ‘Hybrid’ Super Sensitivity SP Detector with the emission range of 493-550 nm. SCRI Renaissance 2200 (SR2200) dye was excited with a Diode 405 nm ‘UV’ laser and images were collected by using PMT SP Detector with the emission 415-470nm. EGFP and SR2200 fluorescence emission was collected with HyD ‘Hybrid’ Super Sensitivity SP Detector (Leica Microsystems) and PMT SP Detector (Leica Microsystems). Z-stacks were created via sequential line scanning. Quantitative image measurements were performed using the Fiji image processing software. Two-dimensional slices from the center of the SAM were chosen based on analysis of Z-stacks to determine the width and height of reporter expression. Spot detection tool of IMARIS software was used to calculate the number of cells in Fig. 5G and H. Nuclei were detected on the base of EGFP, and estimated nuclei diameter values were used for background subtraction.

To observe the GFP signal in pAMO113 seedlings in response to Dex, a 12-megapixel cooled color Nikon DXM-1200c camera and a Nikon Eclipse 80i microscope were used.

### Construction of the mathematical model

We assumed that there is a two-fold symmetry of the SAM. We used 326 points in a quarter ball (half dome) with a radius of 400 length units to represent cells in half of the SAM. The number of cells was estimated from Chen et al. (Chen et al., 2013). In the 3D cell network model, the EPFL ligands are synthesized in two peripheral regions represented by two ‘corners’ regions. We assumed that EPFL diffuses broadly in the SAM and inhibits the expression of both *WUS* and *CLV3* through binding to their receptors which were assumed to be always expressed in each cell (Kosentka et al., 2019; Zhang et al., 2021). Because CLV3 is a diffusive peptide, and WUS is a transcription factor capable of moving between cells (Lenhard and Laux, 2003; Yadav et al., 2011; Daum et al., 2014), we assumed that these molecules are diffusible in the model.

Our model includes WUS-CLV3 negative feedback and its lateral regulator EPFL. In addition, it describes a HAIRY MERISTEM (HAM) signal that originates from the rib zone and inhibits *CLV3* expression in the organizing center (Zhou et al., 2018). The distribution of *HAM* expression is likely established by other signals not considered in the model (Han et al., 2020b). It was shown that a high concentration of WUS can cause *CLV3* downregulation, forming a biphasic regulation of *CLV3* by WUS (Perales et al., 2016; Shimotohno and Scheres, 2019), and this regulation is described in our model. The model also includes a CLV3 independent positive feedback involving WUS. This feedback can be supported by a WUS-cytokinin mutual activation loop: it was previously shown that cytokinin activates *WUS* expression (Gordon et al., 2009; Chickarmane et al., 2012; Wang et al., 2017), whereas WUS derepresses cytokinin signal by inhibiting Type A *ARABIDOPSIS RESPONSE REGULATOR* (ARR) genes which act as inhibitors of cytokinin (To et al., 2004; Leibfried et al., 2005; Shimotohno and Scheres, 2019).

This positive feedback loop may also be supported by other factors (Yadav et al., 2013). Based on these assumptions, the dynamics of six interacting species representing concentrations of regulatory molecules are described with nonlinear ordinary differential equations (ODEs) in each cell (point) of the model (additional spatial constraints are shown in Figure 7B):

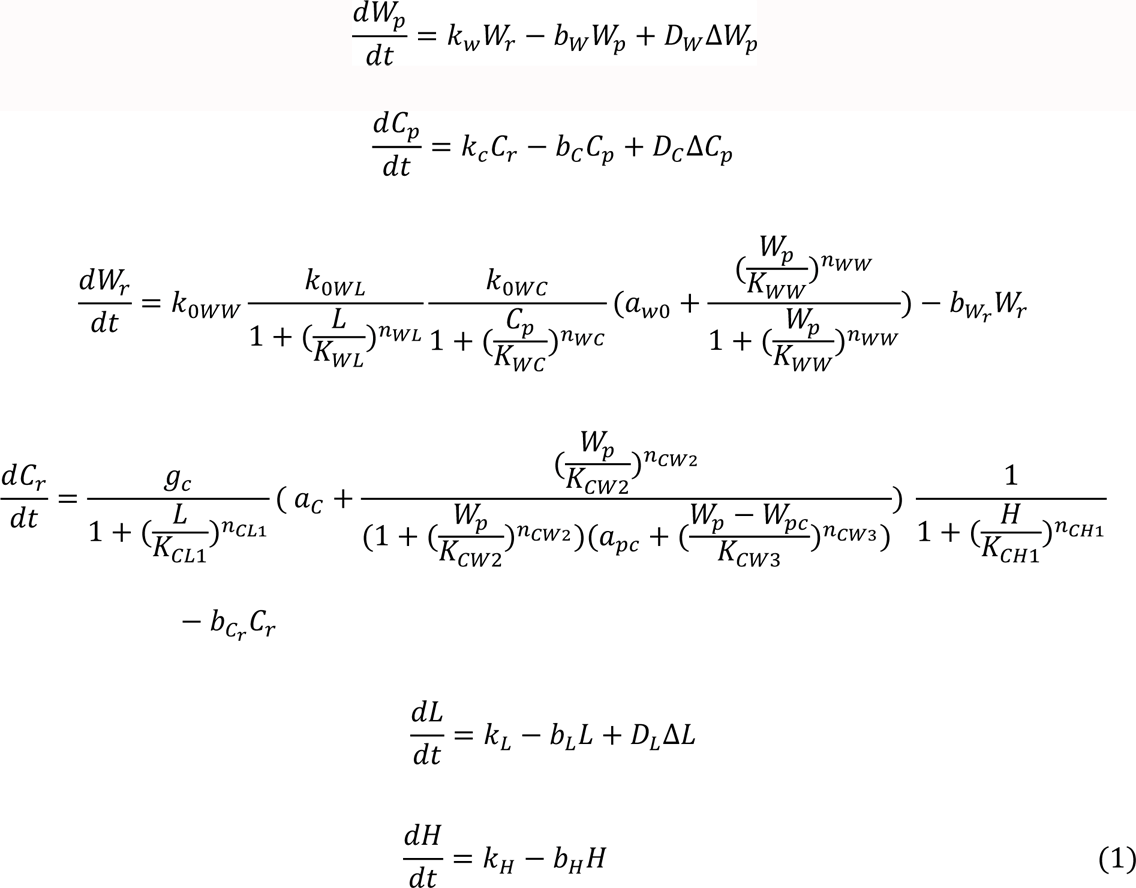

Here, state variables *W*_r_, *W*_p_, *C*_r_, *C*_p_, *L* and *H* represent the concentrations (or strengths) of *WUS* mRNA, WUS protein, *CLV3* mRNA, CLV3 protein, EPFL, and HAM respectively. A full list of parameter descriptions and their numerical values is available in Supplemental Table 3. In the ODEs, *b*_w_ is the production rate constant of *WUS* protein; *b*_wc_ is the degradation rate constant of *WUS* protein; *K*_*Wc*_ is the threshold of inhibition of *WUS* by *CLV*3. *n*_wC_ is the cooperativity of inhibition of *WUS* by *CLV*3; *D*_w_ is the rate constant of passive diffusion-like transport of molecule *WUS protein*; *k*_C_is the production rate constant of *CLV*3 protein. *D*_C_ is the degradation rate constant of CLV3 protein. *D*_C_ is the rate constant of passive diffusion-like transport of molecule CLV3 protein. *k*_Oww_ is the production rate constant of *WUS* mRNA. *k*_OwL_ is the proportion of *WUS* mRNA production rate controlled by EPFL. *K*_wL_ is the threshold of *WUS* inhibition by EPFL. *n*_wL_ represents the cooperativity of regulation of *WUS* by EPFL. *k*_OwC_ is the proportion of *WUS* mRNA production rate controlled by CLV3. *K*_ww_ is the activation threshold of WUS autoactivation. *n*_ww_ is the cooperativity of WUS self-regulation. *K*_wr_ is the degradation rate constant of *WUS* mRNA. *g*_C_ is the production rate constant of *CLV3* mRNA. *K*_CL1_is the threshold of *CLV3* inhibition by EPFL. *n*_CL1_ cooperativity of regulation of *CLV3* by EPFL. *K*_Cw2_ is the threshold of *CLV3* activation by WUS. *n*_Cw2_ is the cooperativity of regulation of *CLV3* by WUS. *a*_pC_ is the constant representing the inversed strength of *CLV3* inhibition by WUS. *K*_Cw3_ is the threshold of *CLV3* inhibition by WUS. *n*_Cw3_represents the cooperativity of negative regulation of *CLV3* by WUS. *K*_CH1_ is the threshold of *CLV3* activation by HAM. *n*_CH1_represents the cooperativity of regulation of *CLV3* by HAM. *K*_Cr_ is the degradation rate constant of *CLV3* mRNA. *k*_L_ is the production rate constant of EPFL. *K*_L_ is the degradation rate constant of EPFL protein. *D*_L_ is the passive diffusion rate constant of EPFL protein. *k*_H_ is the production rate constant of HAM. *K*_H_ is the degradation rate constant of HAM protein; Δ is the Laplace operator describing gradients of concentrations, which govern passive diffusion-like transport; Δ*W*_p_, Δ*C*_p_, Δ*L* have a unit of concentration per unit area. *D*_w_, *D*_C_, *D*_L_ were adjusted by multiplying with a scaling factor 〈*l*〉/*l*, where *l* represents the distance between the centers of the two cells (Delile et al., 2017); and neighboring cells are defined as cells that are located within a radius of 100 length units (∼10 *μm*). We neglected the subcellular geometry of the cells, their contact areas, and the influence of mechanics in this study (the effected contact area for WUS transport cannot be directly inferred from total contact area of plasma membrane). The movements of EPFL, CLV3 and WUS are responsible for the intercellular communication in the model. We used a Hill function to describe nonlinearity in the gene regulation. Previous models of the SAM and other complex systems have used similar nonlinear functions (Nikolaev et al., 2007; Fujita et al., 2011; Gruel et al., 2018; Ye et al., 2019; Liu et al., 2020). *a*_C_ is a constant used to perturb the negative feedback regulation. *a*_wO_is a constant used to perturb the auto-positive feedback regulation. Because the absolute concentrations of these molecules have not been measured experimentally, we used an arbitrary unit (a.u.) to describe concentration (or strength) of each molecule. We used a no-flux boundary condition for the model, as in other published SAM models (Zhou et al., 2018; Liu et al., 2020).

We fitted the parameters to known patterning phenotypes of the SAM under normal and genetically perturbed conditions. The *erf* mutant was represented by setting the EPFL production rate to 0. The *clv3* mutant was represented by setting the CLV3 protein production rate to 0.

Each regulation-specific mutant was modeled by setting the inhibition threshold to large number (1000). Because only qualitative information is available from the experimental data, we performed the fitting manually. To perform a simulation for a SAM system, we solved the system of ODEs numerically using the Tellurium package (Choi et al., 2018). The initial concentrations for all variables were set to zero. For all our analyses, steady state solutions (at time unit 98) were used to determine the patterning of the SAM. For visualization of gene expression levels, expression values were normalized to [0, 1] by dividing each value by the maximum level of the molecule across all conditions.

## Acknowledgments

We thank Jaydeep Kolape at Advanced Microscopy and Imaging Center, UTK for technical assistance. We thank Guangzhong Lin and Jijie Chai for sharing with us EPFL6 protein and Ian Moore for sharing with us GR-LhG4:T35S, pBIN-LhGR-N, and pH-TOP plasmids.

## Author contributions

E.D.S and T.H conceived the research plan and supervised the experiments; M.U., R.U.C., Z. L., A.O., D.D., L. Z., H.B., T.H., and E.D.S performed the experiments and analyzed the data; E.D.S, T.H., and M.U wrote the article with contributions of all the authors; E.D.S. agrees to serve as the author responsible for contact and ensures communication.

## Data and materials availability

All data are available in the main text or the supplementary materials.

## This article contains supporting information

Supplemental Figure 1 Inter- and intragroup sample variability of transcriptomics data

Supplemental Figure 2 The structure of T-DNA insert in the pAMO113 vector.

Supplemental Figure 3 An example of *WUS* expression in a SAM observed at different angles.

Supplemental Table 1 Transcriptome dataset

Supplemental Table 2 Primers used in this study.

Supplemental Table 3 Parameter values of the wild-type SAM mathematical model.

Supplemental Video 1 WUSp: H2B-GFP expression in 3DPG the wild type SAM

Supplemental Video 2 WUSp: H2B-GFP expression in 3DPG *er erl1 erl2*

Supplemental Video 3 WUSp: H2B-GFP expression in 3DPG clv3

